# BRCA2-deficiency Causes Global Transcriptomic Alterations in Endothelial Cells

**DOI:** 10.1101/2025.09.14.676113

**Authors:** Sepideh Nikfarjam, Aman Singh, Shuhan Bu, Hien C. Nguyen, Krishna K. Singh

**Author notes:** To whom correspondence should be addressed: Krishna K. Singh, Department of Medical Biophysics, Schulich School of Medicine & Dentistry, Western University, 1151 Richmond St. N., London, Ontario N6A 5C1, x85683 (Lab). **Ethical approval statement:** Not applied. **Author contribution statement:** K. K. S. conceptualization; S. N., A.S., S. B., and K. K. S. methodology and investigation; S. N., H.N., and K. K. S. data curation and formal analysis; S. N. writing—original draft; K. K. S., and A. S. writing—review & editing; S. N. visualization; S. N., A.S. and K. K. S. validation; K. K. S. supervision; K. K. S. resources; K. K. S. project administration; K. K. S. funding acquisition. **Data availability statement:** Not applied. **Consent statement:** Not applied.

## Abstract

This study aims to explore the alterations in gene expression following *BRCA2* loss in endothelial cells and to investigate the potential contribution of *BRCA2* in regulating the endothelial protein-coding transcriptome. Cultured human umbilical vein endothelial cells (HUVECs) were transfected with siBRCA2 or non-targeting scrambled RNA as control. Total RNA was extracted and used for RNA sequencing following purification and quality check. Transcription profiles and differentially expressed genes (DEGs) were identified. Enrichment of functions and signaling pathways analysis were performed based on Gene Ontology (GO) and the Kyoto Encyclopedia of Genes and Genomes (KEGG) database. In total, 3196 DEGs with (cut-off 1.5-fold) were identified, of which 1852 genes were upregulated, and 1344 genes were downregulated in *BRCA2*-deficient HUVECs. *Cell cycle exit and neuronal differentiation 1* (CEND1) and *mannose receptor C-type 1* (MRC1) were the most significantly up- and downregulated genes, respectively, in BRCA2-deficient endothelial cells. The expression of top DEGs were further validated by RT-qPCR in HUVECs and in human coronary artery endothelial cells. The GO and KEGG analysis suggest that cell adhesion molecules and cytokine-cytokine receptor interaction may play an important role in BRCA2-deficient endothelial cells. Overall, these findings contribute to a greater understanding of the mechanisms involved in BRCA2-mediated endothelial function.

## Introduction

Maintaining genomic stability is a vital priority which supports normal cell survival and proliferation [1]. Any aberrant genetic alterations can interfere with regular cellular processes and functions [2]. Accordingly, cells have evolved a variety of signaling pathways, collectively known as DNA damage repair (DDR) signaling, to evaluate DNA metabolism, hamper the accumulation of DNA damage, and maintain genomic stability [3]. Persistent DDR and accumulated DNA damage along with dysfunctional DDR proteins can lead to programmed cell death or carcinogenesis. Interestingly, a growing amount of clinical and preclinical evidence has revealed the presence of DNA damage and activation of DDR signaling in cardiovascular diseases (CVDs) [4-6]. There are remarkable commonalities between cancer and CVDs including common risk factors and intracellular pathways for disease development and progression. Many of the “hallmarks of cancer”, such as inflammation, genomic instability, cellular proliferation, cell death resistance, and angiogenesis represent pathophysiologic processes common to both cancer and CVDs [7].

Among various cellular components orchestrating DDR, the breast cancer susceptibility genes, *BRCA1* and *BRCA2 (BRCA1/2)*, play critical roles in maintaining genome integrity by participating in homology-directed repair of DNA double-strand breaks (DSBs) as well as regulation of transcription and cell cycle progression [8]. Loss of BRCA1/2 function results in defective DDR that, if left unchecked, leads to accumulation of damaged DNA, resulting in carcinogenesis or p53-mediated apoptosis [9]. More importantly, recent studies reported a critical role for *BRCA1/2* genes in cardiovascular dysfunction. Loss- and gain-of BRCA1 function exacerbated and protected against atherosclerosis-induced endothelial dysfunction by modulating DDR pathways in endothelial cells [10]. In laboratory studies, mice harboring cell-specific *BRCA1* mutations were more susceptible to cardiovascular dysfunction such as endothelial dysfunction in atherosclerosis and cardiomyopathy secondary to myocardial infarction and genotoxic drug treatment [10-14]. BRCA2 is a relatively understudied protein in the context of CVDs. In addition to DDR, BRCA2 is associated with regulation of lipid metabolism and inflammation in humans [15-18]. Interestingly, BRCA2 expression is modulated in a context-dependent manner. While stress in the form of exposure to oxidized low-density lipoprotein decreased endothelial expression of BRCA2 [19], and high glucose enhanced BRCA2 expression in endothelial cells [20]. BRCA1 and BRCA2 play non-redundant roles in different stages of genome protection, and lead to different cancer predisposition rates, frequency, and lifetime risk. In contrast to BRCA1, BRCA2 is also associated with lipid regulation, inflammation and elevated risk of coronary artery disease [19]. Moreover, the BRCA2 gene resides on human chromosome 13q12.3, a region highly linked to CVD [19]. Therefore, loss of function studies on endothelial cell-specific BRCA2 mutation will provide novel information related to cardiovascular health and disease, and identifying molecular targets and mechanisms linked to BRCA2-deficiency in cells of the cardiovascular system would provide novel information regarding the pathogenesis of CVDs in BRCA2 mutations.

In this study, we report for the first time a potential role for BRCA2 in regulating the endothelial transcriptome. We silenced *BRCA2* in cultured endothelial cells, and the effect of loss of BRCA2 function on protein-coding transcriptome was assessed by RNA sequencing. Interestingly, our findings revealed a total of 3196 differentially expressed genes between the BRCA2-deficient HUVECs and the scrambled control-transfected cells. This result indicates the necessity for further investigation on the role of BRCA2-associated mechanisms in endothelial cells as well as other cells of the cardiovascular system.

## Material and Methods

### Cell culture

human umbilical vein endothelial cells (HUVECs, Lonza, mixed) and human coronary artery endothelial cells (HCAECs, Lonza, mixed) were cultured in endothelial cell growth medium (EGM™-2 Bulletkit™; Lonza) supplemented with growth factors, 5% fetal bovine serum (FBS), and antibiotics at 37°C and 5% CO_2_. Passage 4 HUVECs and passage 5 HCAECs were used for downstream experiments.

### BRCA2 silencing

Small interfering RNA (siRNA) against *BRCA2* transcript (Dharmacon™, #A-003462-13-0005) as well as non-targeting scrambled control RNA (Dharmacon™, #2575450) were used. HUVECs were transfected with siBRAC2 or scrambled control using the Lipofectamine™ 3000 transfection reagent (Invitrogen) according to the manufacturer’s protocol as previously described [19]. Briefly, 2×10^5^ HUVECs or HCAECs were seeded in 6-well cell culture plates and transfected with 5 nM siBRCA2 or scrambled control (n=4/group) and incubated in endothelial cell growth medium. All experiments were performed 48 h post-transfection.

### RNA extraction and purification

HUVEC or HCAEC cultures were washed once with ice-cold PBS. TRIzol™ (Invitrogen) was added and incubated for 10 min at room temperature, and total RNA was extracted as instructed by the manufacturer. Following extraction, RNA was further purified by mixing with 0.1 volumes of 3M sodium acetate and 2.5-3 volumes ice cold 100% ethanol and subsequent precipitation overnight at −20°C. The mixture was then centrifuged at 13000 rpm, at 4°C for 30 minutes, and the pellet was washed twice with 500 μL ice cold 75% ethanol at 4°C for 10 mins each time. The pellet was air dried and resuspend in 20 μL of nuclease-free water.

### RNA sequencing

Eight samples were sequenced at the London Regional Genomics Centre (Robarts Research Institute, London, Ontario, Canada; http://www.lrgc.ca) using the Illumina NextSeq 500 (Illumina Inc., San Diego, CA). Total RNA quality was evaluated using the NanoDrop ND-1000 spectrophotometer (Thermo Fisher Scientific, Waltham, MA), the Agilent 2100 Bioanalyzer (Agilent Technologies Inc., Palo Alto, CA), and the RNA 6000 Nano kit (Caliper Life Sciences, Mountain View, CA) (*Supplementary Figure 1*). Samples with an RNA integrity number (RIN) > 9.0 were used for downstream analysis (n=3/group). The samples were then processed using Total RNA-seq (H/M/R) Library Prep Kit for Illumina (Vazyme, Nanjing, China). Briefly, samples were rRNA depleted, fragmented, and utilized for cDNA synthesis and PCR amplification with indexed primers to permit equimolar pooling of samples into one library. The pooled library size distribution was assessed on an Agilent High Sensitivity DNA Bioanalyzer chip and quantitated using the Qubit 2.0 Fluorometer (Thermo Fisher Scientific, Waltham, MA). The library was sequenced on the Illumina NextSeq 500 as single end runs, 1 × 76 bp, using High Output v2 kits (75 cycles). Fastq data files were analyzed using Partek Flow (St. Louis, MO). After importation, the data were aligned to the *Homo sapiens* genome hg19 using STAR 2.7.3a and annotated using RefSeq Transcripts. Features with more than 18 reads were normalized using Trimmed Mean of M-values (TMM) (https://doi.org/10.1186/gb-2010-11-3-r25) followed by adding 0.0001. Any batch effect due to the run date was removed using Partek Flow’s Remove batch effect tool based on the results of the principal component analysis (PCA) plot. The gene-specific analysis (GSA) function of Partek Flow was then used to determine differential gene expression in BRCA2-silenced versus scramble control-transfected HUVECs using Akaike Information Criteria corrected – a repeated measure analysis using mixed models’ methodology. The filtered gene list (fold change ≥ 1.5 and FDR ≤ 0.05) was then used for downstream analyses.

### Gene Ontology (GO), Kyoto Encyclopedia of Genes and Genomes (KEGG), and Metascape enrichment analyses

To identify enriched biological themes over-represented in BRCA2-deficient HUVECs, we identified the upregulated genes with the cut-off of fold-change ≥ 1.5 and FDR ≤ 0.05 and then analyzed GO terms and KEGG pathways enriched in these differentially expressed gene (DEG) sets. For functional enrichment analysis, all DEGs were mapped to terms in the GO databases, and then significantly enriched GO terms were searched for among the DEGs using *p* < 0.05 as the threshold. GO term analysis was classified into three subgroups, namely biological processes (BP), cellular component (CC) and molecular function (MF). All DEGs were mapped to the Reactome (https://maayanlab.cloud/Enrichr/) and KEGG (https://www.genome.jp/kegg/) databases and searched for significantly enriched pathway terms at *p*<0.05 level. The filtered gene list was then submitted to Metascape (https://metascape.org/gp/index.html#/main/step1) using Express Analysis function of *Homo sapiens* gene IDs. A subset of enriched terms was selected and rendered as a network plot, where terms with a similarity > 0.3 were connected by edges. Terms with the lowest *p*-values from each of the 20 clusters were selected, with the constraint that there are no more than 15 terms per cluster and no more than 250 terms in total. The network was visualized using Metascape, where each node represented an enriched term and was colored first by cluster ID and then by *p*-value.

### Reverse transcription quantitative polymerase chain reaction (RT-qPCR)

The Quantitect kit (Qiagen) was utilized to synthesize complementary DNA, which was then subjected to qPCR using the ABI ViiA 7 Real-Time PCR System (Applied Biosystems). For PCR, Power SYBR™ Green PCR Master Mix (Applied Biosystems) were mixed with forward and reverse primers for human *GAPDH, BRCA2, Cell Cycle Exit and Neuronal Differentiation 1 (CEND1), EGF-Like Repeats and Discoidin Domains 3 (EDIL3), Potassium Voltage-Gated Channel Subfamily A Regulatory Beta Subunit 1 (KCNAB1), Janus Kinase and Microtubule Interacting Protein 2 (JAKMIP2), Cytoplasmic Polyadenylation Element Binding Protein 2 (CPEB2), Coiled-Coil Domain Containing 136 (CCDC136), Family with Sequence Similarity 83 Member B (FAM83B), F-Box Protein 10 (FBXO10), Six Transmembrane Epithelial Antigen of the Prostate 3 (STEAP3), Interleukin 10 Receptor Subunit Beta (IL10RB), CXADR-like membrane protein (CLMP), Mannose Receptor C-Type 1 (MRC1), Retinoic Acid Early Transcript 1E (RAET1E), Cytidine Monophosphate Kinase 1 (CMPK1), Leucine-Rich Repeat Transmembrane Neuronal 2 (LRRTM2), N-Acetyltransferase 8B (NAT8B) and G protein-coupled receptor 1 (GPR1)* (*Supplementary Table 1*) in accordance with the manufacturer’s instructions. The primer sequences were obtained from a PCR primer database (https://pga.mgh.harvard.edu/primerbank/) for every gene. A serial dilution of cDNA was performed and then primer with efficiency >90% were used. qPCR was performed for n=3 in triplicates. Gene expression was analyzed using the ΔΔCt method. For each gene, Ct values were first normalized to the corresponding housekeeping gene (ΔCt), and then ΔCt values from siBRCA2-transfected cells were compared to those of the scrambled control-transfected group to calculate ΔΔCt.

### Statistical analysis

For RT-qPCR, statistical comparisons between groups were performed using Student’s *t*-test on ΔΔCt values, which are inherently log_2_-transformed and commonly used for parametric analysis. Data were normally distributed with equal variances. GraphPad Prism version 10.2.3.4.403 (GraphPad Software, San Diego, CA, USA) was used to perform Student’s *t*-test, and *p* < 0.05 was considered statistically significant.

## Results

### Identification of differentially expressed transcripts in BRCA2-deficient endothelial cells

To reveal the molecular targets associated with loss of BRCA2 function in endothelial cells, we profiled the protein-coding RNAs of BRCA2-silenced HUVECs and compared this to scrambled control-transfected cells. The principal component analysis (PCA) plot **(Figure 1)** shows clear separation of total RNA samples of *BRCA2*-silenced HUVECs from the control cells along the principal components (PCs) with the overall variance of 97.87%. The separation along PC1 accounts for most of the variance in the data (74.16%) **(Figure 1)**. A DEG analysis was performed to identify gene expression changes between the two cell groups. A total of 3196 DEGs (±1.5 fold-change and adjusted *p*-value < 0.05) were detected between *BRCA2*-silenced and control cDNA libraries, of which 1852 genes were significantly upregulated, and 1344 genes were significantly downregulated upon BRCA2-deficiency **(Figure 2**, *Supplementary Table 2*). The expression of 17457 genes remained unaltered **(Figure 2)**. The top upregulated DEGs in *BRCA2*-silenced HUVECs included *CEND1, KCNAB1, JAKMIP2, CCDC136, FAM83B, STEAP3, IL10RB, CLMP*, and *EDIL3* **(Table 1)**. The top downregulated DEGs in *BRCA2* knockdown HUVECs were *MRC1, RAET1E, DNASE1L3, CMPK1, LRRTM2, NAT8B* and *GPR1* **(Table 2)**. The heat map generated using z-score revealed significant alterations in gene expression of BRCA2-deficient HUVECs compared to control cells **(Figure 3)**.

**Figure 1.**
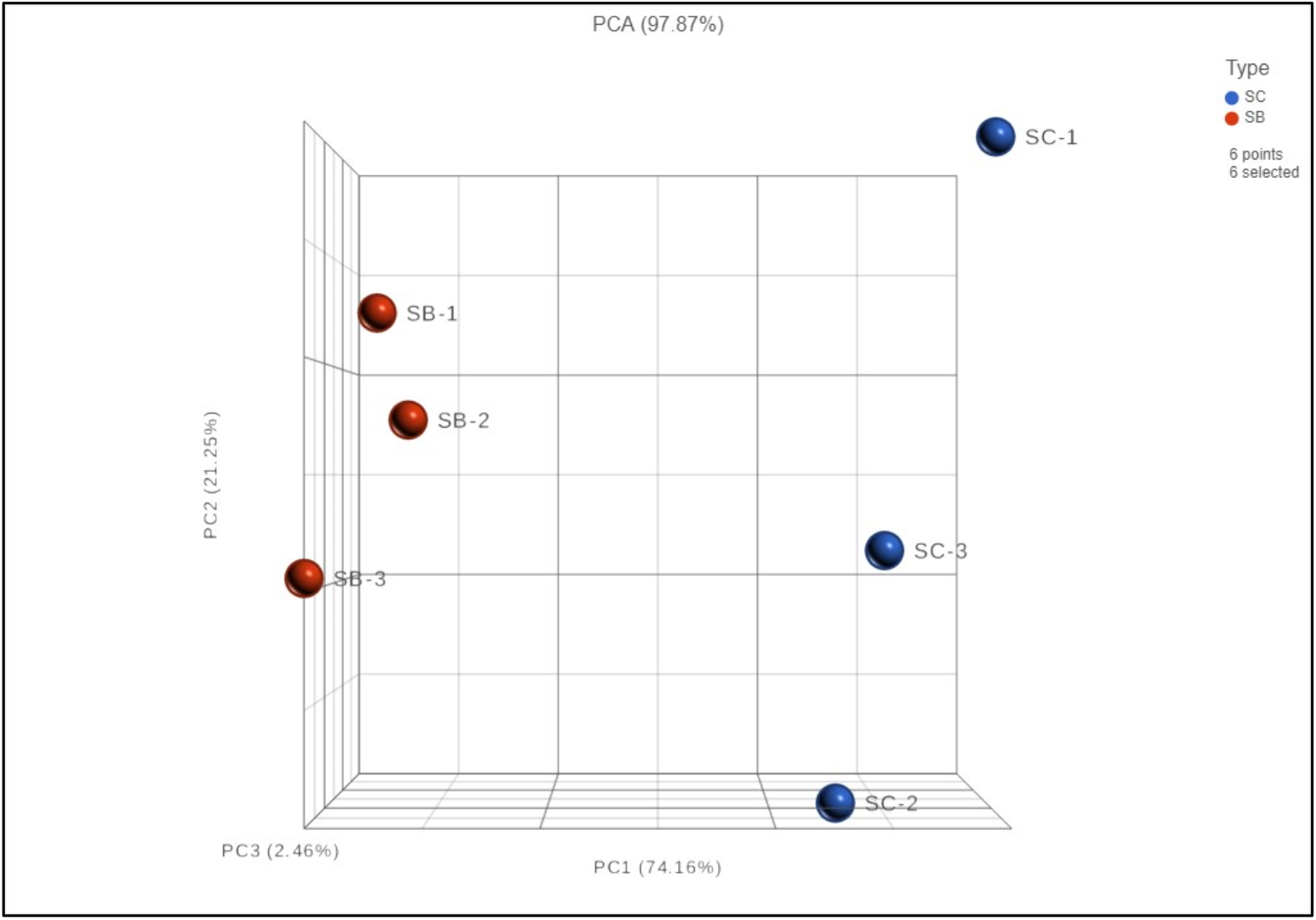
Principal component analysis plot of total RNA samples derived from *BRCA2*-silenced and scrambled control RNA-transfected human umbilical vein endothelial cells. The plot shows the first three principal components (PC1, PC2, and PC3) and the overall variance (PCA: 97.87%) with their respective explained variance percentages. Data points represent siBRCA2-transfected (SB) and scrambled control RNA-transfected HUVECs (SC), labeled in red and blue, respectively.

**Table 1.** Top 10 upregulated differentially expressed genes between *BRCA2*-silenced and scrambled control RNA-transfected HUVECs and HCAECs.

| RNAseq (HUVECs) |  |  | Validation RT-qPCR (HUVECs) |  | Validation RT-qPCR (HCAECs) |  |
| --- | --- | --- | --- | --- | --- | --- |
| Gene | Fold-change | *p*-value | Fold-change (Mean ± SD) | *p*-value | Fold-change (Mean ± SD) | *p*-value |
| *CEND1* | 50.53 | 8.0621E-27 | 38.20±4.47 | 4.2552E-08 | 2.02±0.66 | 0.016 |
| *KCNAB1* | 18.59 | 5.295E-101 | 41.43±4.75 | 3.6E-08 | 6.45±3.03 | 0.015 |
| *JAKMIP2* | 18.03 | 1.4393E-13 | 11.57±2.22 | 1.4889E-06 | 1.80±0.70 | 0.024 |
| *CCDC136* | 13.82 | 3.0765E-39 | 8.28±0.92 | 3.5E-10 | 2.20±0.80 | 0.014 |
| *FAM83B* | 11.62 | 3.6088E-07 | 3.85±1.60 | 0.00163 | 2.08±0.71 | 0.029 |
| *STEAP3* | 11.60 | 0.00E+0 | 11.63±0.94 | 1.831E-09 | 1.92±0.22 | 0.002 |
| *IL10RB* | 11.32 | 3.0367E-21 | 4.58±0.56 | 3.5398E-08 | 1.96±0.858 | 0.039 |
| *CLMP* | 11.15 | 3.6818E-41 | 14.11±2.12 | 4.8558E-08 | 2.29±0.66 | 0.010 |
| *EDIL3* | 10.43 | 1.806E-177 | 8.44±0.68 | 2.5E-10 | 12.45±5.05 | 0.002 |

**Table 2.**
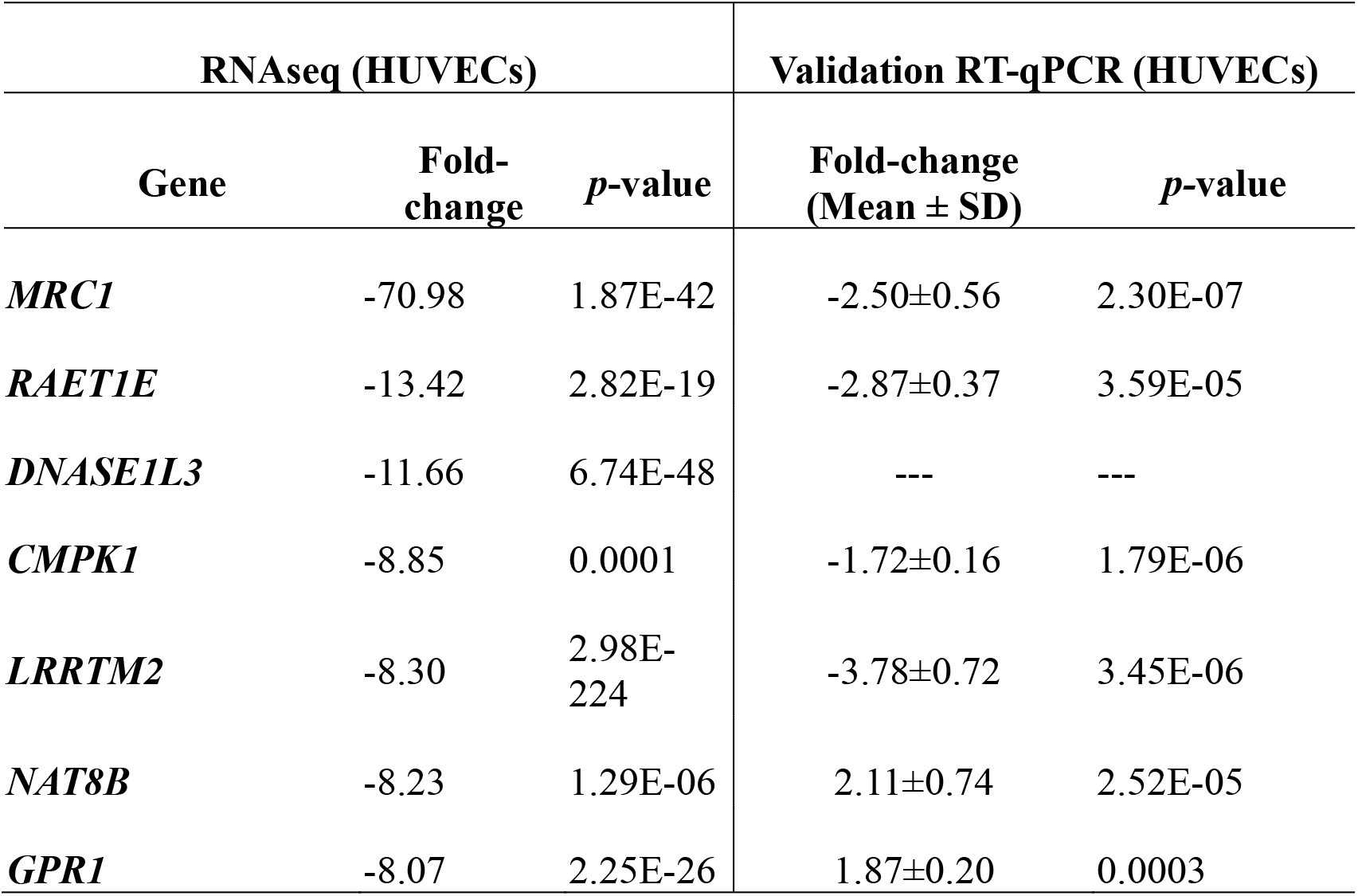
Top 7 downregulated differentially expressed genes between *BRCA2*-silenced and scrambled control RNA-transfected HUVECs.

**Figure 2.**
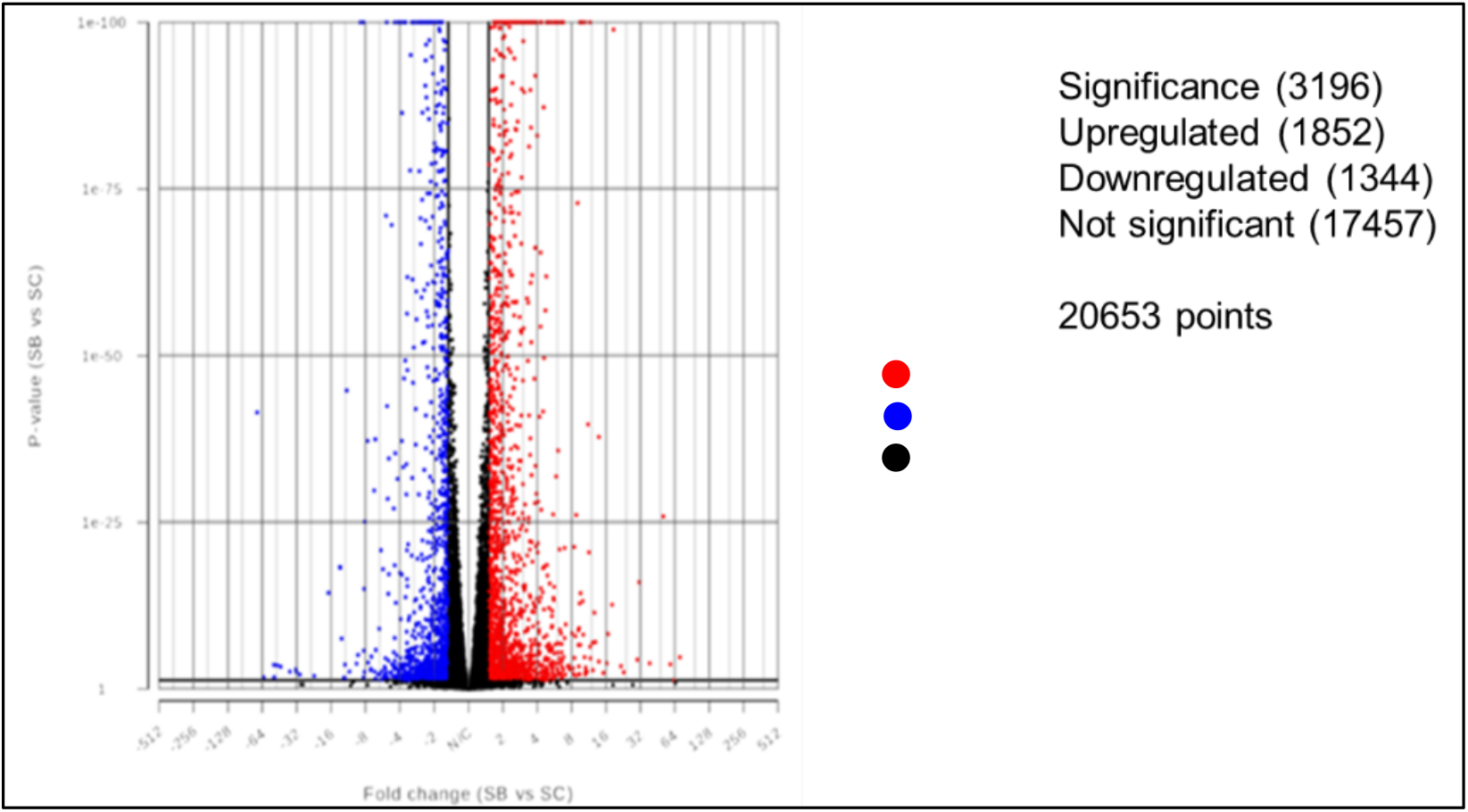
Volcano plot of differentially expressed genes between *BRCA2*-silenced and scrambled control RNA-transfected human umbilical vein endothelial cells. The x-axis is the log2 scale of the fold-change (log2(fold-change)) of gene expression in *BRCA2*-silenced (SB) and scrambled control RNA-transfected (SC) HUVECs. Positive values are indicative of gene upregulation, and negative values are indicative of gene downregulation. The y-axis is the minus log10 scale of the adjusted *p* values (-log10(*p* value)), which represents the significance level of expression difference. The red dots indicate significantly upregulated genes with at least 1.5-fold-change, and the blue dots indicate significantly downregulated genes with at least 1.5 fold-change. Black dots indicate that there is no statistically significant difference in gene expression. The greater ordinate value corresponding to each point indicates greater difference in gene expression corresponding to that point. Similarly, the greater value of the abscissa shows greater difference in gene expression corresponding to that point.

**Figure 3.**
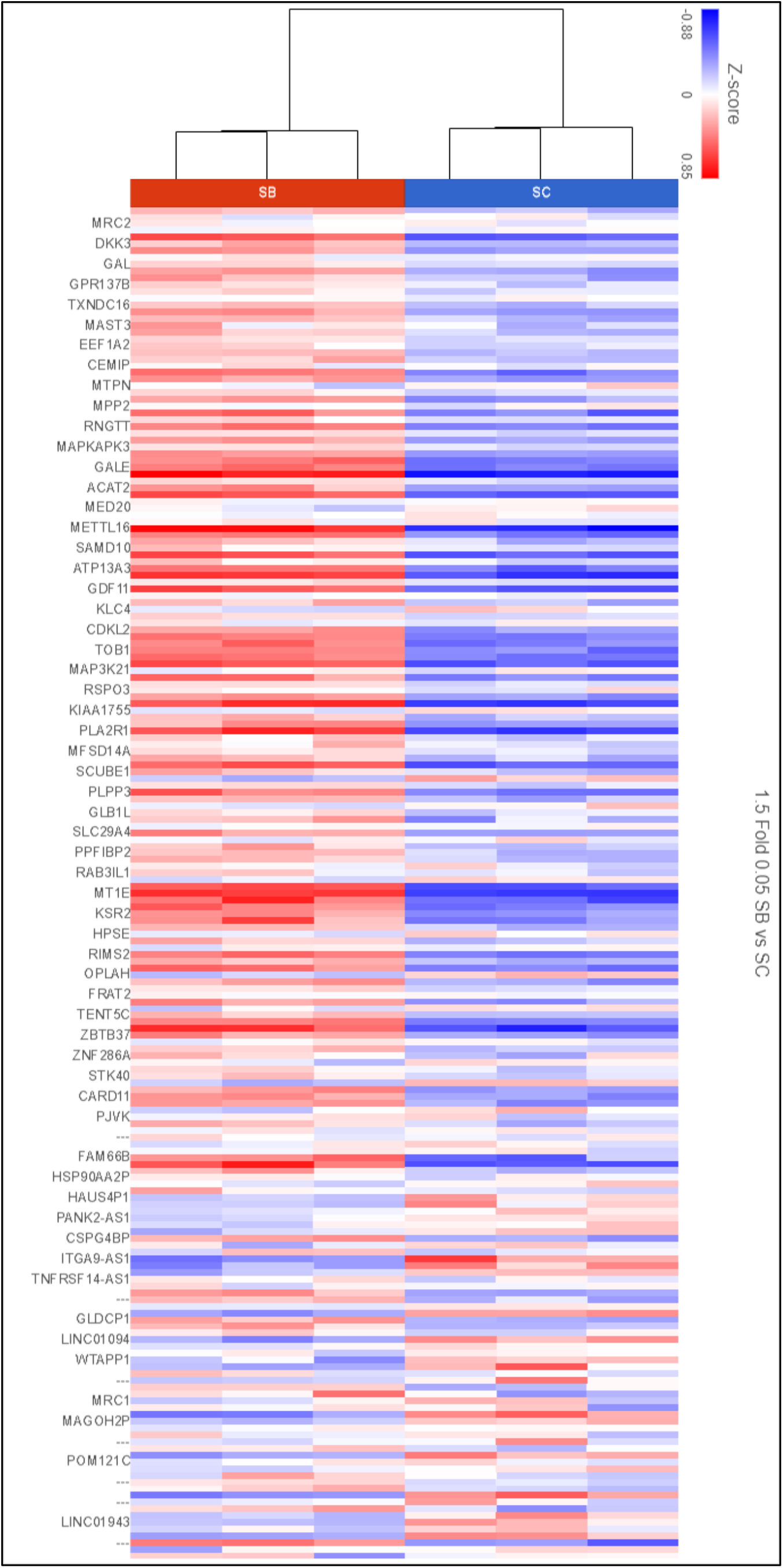
Heat map of differentially expressed genes between *BRCA2*-silenced and scrambled control RNA-transfected human umbilical vein endothelial cells. Hierarchical clustering dendrogram of gene expression between siBRCA2-transfected (SB) and scrambled control RNA-transfected (SC) HUVECs. The horizontal axis at the top represents the name of samples. The vertical axis on the left represents the name of genes. The red and blue stripes represent up- and downregulated DEGs, respectively. Darker red indicates a stronger upregulation in expression and the darker blue indicates a stronger downregulation in expression.

### Functional annotation and classification of differentially expressed genes in BRCA2-silenced HUVECs

Gene Ontology (GO) analyses were carried out to examine three different aspects, namely Biological Processes (BP), Cellular Component (CC), and Molecular Function (MF), reflecting the dynamic alteration processes in BRCA2 deficiency in HUVECs (*Supplementary Table 3*). The top ten GO terms in the enrichment analysis are presented in **Figure 4**. The most enriched BP terms include axon guidance, secondary alcohol biosynthetic process, cholesterol biosynthetic process, regulation of cell migration, positive regulation of vasculature development, regulation of smooth muscle cell migration, positive regulation of cell migration and angiogenesis, and positive regulation of endothelial cell proliferation **(Figure 4A)**. The most enriched CC terms were external side of apical plasma membrane, caveola, plasma membrane raft, collagen-containing extracellular matrix, cytoskeleton of presynaptic active zone, cortical cytoskeleton, platelet dense tubular network, membrane raft, exocytic vesicle, and platelet alpha granule **(Figure 4B)**. The most enriched MF terms were acetylgalactosaminyltransferase activity, platelet-derived growth factor binding and receptor binding, carbohydrate proton symporter activity, semaphorin receptor activity, phosphatase activity, muscle alpha-actinin binding, semaphorin receptor binding, alpha-actinin binding, and UDP-glycosyltransefase activity **(Figure 4C)**.

**Figure 4.**
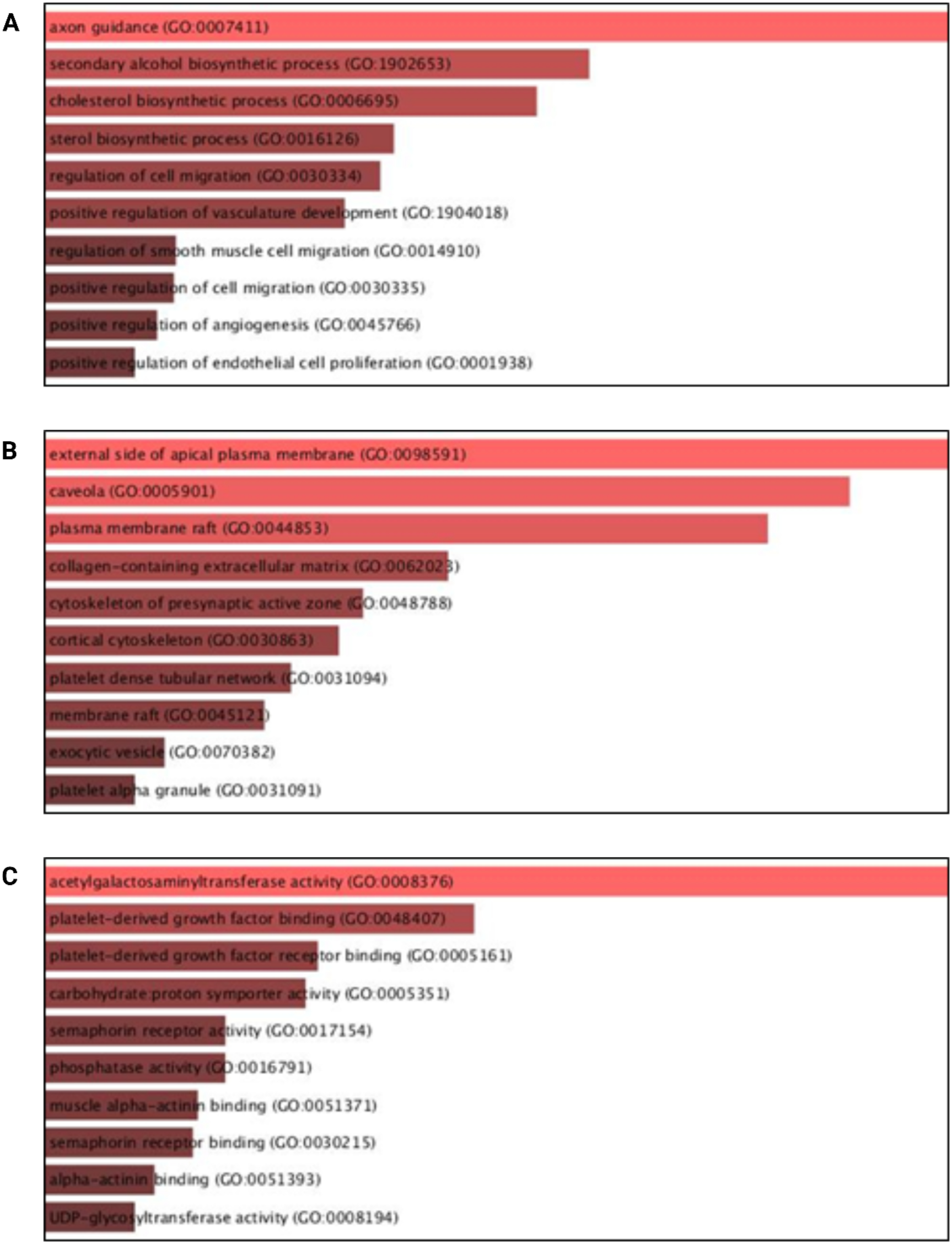
Significantly enriched Gene Ontology (GO) terms between *BRCA2*-silenced and scrambled control RNA-transfected human umbilical vein endothelial cells. The list of 3196 differentially expressed genes was submitted to GO database, and significantly enriched terms with false discovery rate (FDR) < 0.05 were screened out. **(A)** The top ten biological process (BP) terms in the enrichment analysis. **(B)** The top 10 cellular component (CC) terms in the enrichment analysis. **(C)** The top 20 molecular function (MF) terms in the enrichment analysis. UDP: uridine diphosphate.

Next, a KEGG pathway enrichment analysis was conducted to explore the most significantly enriched pathways among the DEGs. A total of 3196 genes were mapped into signaling pathways, and significantly enriched KEGG pathways with false discovery rate (FDR) < 0.05 were screened out **(Figure 5A**, *Supplementary Table 4*). The ranking was according to the enrichment score, *p-* value, and FDR. KEGG biological pathway enrichment analysis found that AGE-RAGE (advanced glycation endproducts – receptor of advanced glycation endproducts) signaling pathway in diabetic complications, focal adhesion, extracellular matrix (ECM)-receptor interaction, cGMP-PKG (cyclic guanosine monophosphate-dependent protein kinase G) signaling pathway, cell adhesion molecules, and cellular senescence were the most important ones among the 325 pathways according to the enrichment score. Functional enrichment analysis performed via Reactome, suggested that most of the significantly DEGs (ranked by *p*-value) were involved in extracellular matrix organization, cholesterol biosynthesis, molecules associated with elastic fibers, non-integrin membrane-ECM interactions, regulation of cholesterol biosynthesis by SPREP, elastic fiber formation, and signaling by TGFB family members (https://maayanlab.cloud/Enrichr/enrich?dataset=70ab0e36faccec34b60841c8c46029e1) **(Figure 5B)**. These findings were independently explored using further functional analysis with Metascape **(Figure 6)**. The list of DEGs was submitted to Metascape using the Express Analysis function of *Homo sapiens* gene IDs. A subset of enriched terms was selected and rendered as a network plot, where terms with a similarity > 0.3 were connected by edges. Each node represents an enriched term and is colored first by its cluster ID **(Figure 6A)** and then by its *p*-value **(Figure 6B)**. These findings indicate that ECM organization, cellular component morphogenesis, and cell migration are highly impacted upon *BRCA2* knockdown in HUVECs. The enrichment of vascular and nervous system development-related terms suggests broader developmental disruptions, potentially implicating BRCA2 in angiogenesis and neurovascular signaling. The inclusion of metabolism of lipids and actin cytoskeleton organization may point to cellular stress responses or cytoskeletal remodeling under BRCA2 loss. Overall, highly connected central clusters shown in Figure 6 suggest coordinated regulation of pathways, not isolated effects.

**Figure 5.**
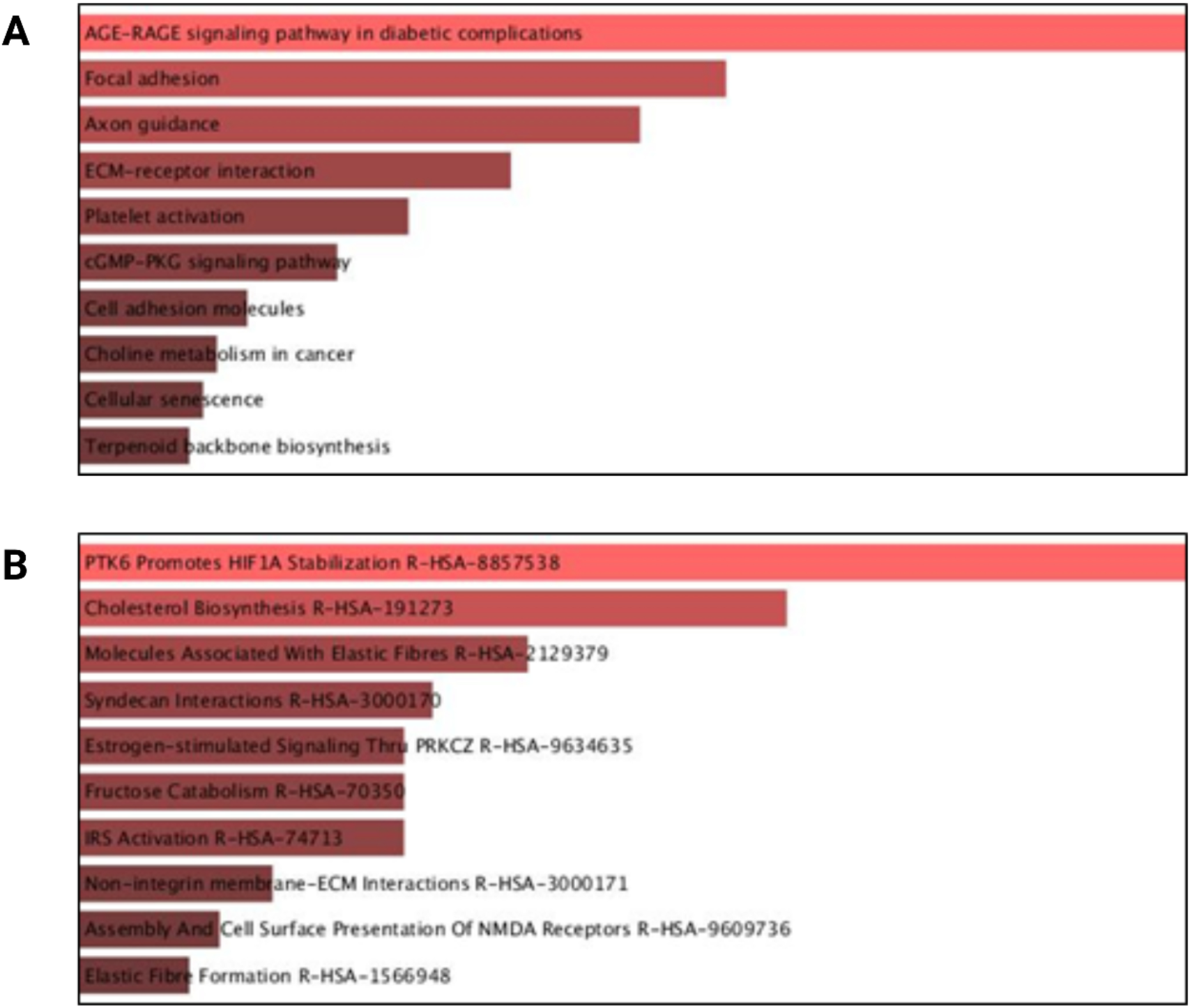
Top enriched Kyoto Encyclopedia of Genes and Genomes (KEGG) and Reactome pathway terms of differentially expressed genes in BRCA2-deficicent human umbilical vein endothelial cells. The list of 3196 differentially expressed genes was submitted to KEGG and Reactome databases for pathway enrichment analysis. **(A)** The top 10 Significantly enriched KEGG pathways were screened out based on their enrichment score, *p* value, and false discovery rate (FDR). **(B)** The top 10 Significantly enriched Reactome pathways with FDR < 0.05 were screened out. AGE-RAGE: advanced glycation endproducts – receptor for advanced glycation endproducts; ECM: extracellular matrix; cGMP-PKG: cyclic guanosine monophosphate-dependent protein kinase G. HIFA1: hypoxia-inducible factor 1 subunit alpha; IRS: insulin receptor signaling. NMDA: N-methyl-D-aspartate; PTK6: tyrosine-protein kinase 6.

**Figure 6.**
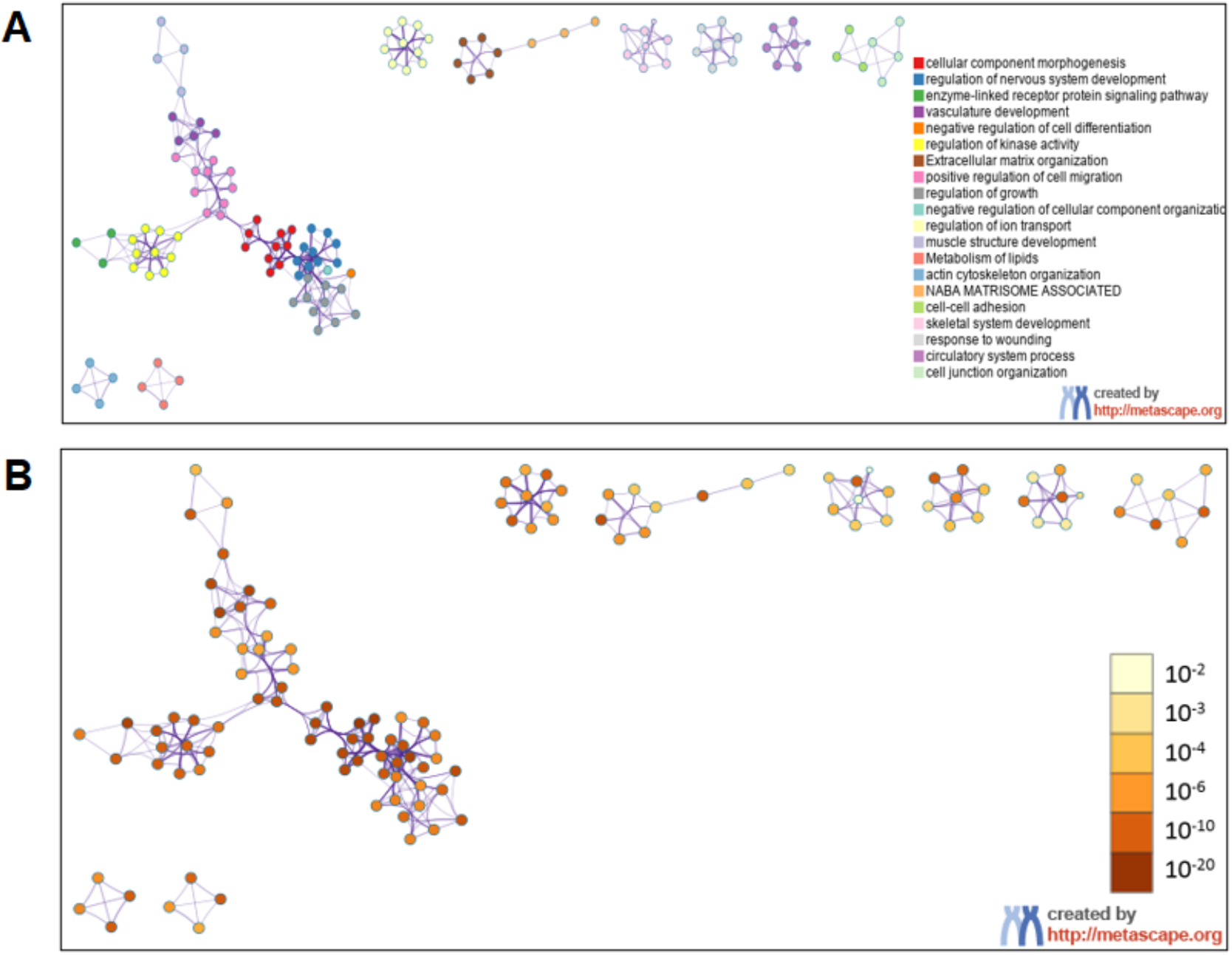
Metascape enrichment network visualization presenting the intra-cluster and inter-cluster similarities of enriched terms in BRCA2-deficient human umbilical vein endothelial cells. The list of 3196 differentially expressed genes was submitted to Metascape and analyzed using the Express Analysis function of *Homo sapiens* gene IDs. A subset of enriched terms was selected and rendered as a network plot, where terms with a similarity > 0.3 were connected by edges. Cluster annotations are shown in color code. Each node represents an enriched term and colored first by its cluster ID **(A**) and then by its *p*-value **(B)**.

### RT-qPCR validation of RNA sequencing results

RT-qPCR validation was performed for the top up- and downregulated DEGs to assess the accuracy of the RNA sequencing data **(Tables 1 and 2)**. Our validation qPCR data in HUVECs confirmed upregulation of top DEGs (*CEND1, JAKMIP2, CCDC136, FAM83B, STEAP3, IL10RB*, and *EDIL3)* identified in the RNA-seq analysis **(Table 1)**. RNA sequencing often detects a wider range of gene expression changes and can sometimes show larger fold changes due to its high sensitivity and broad dynamic range, that can capture more subtle expression changes across different transcript isoforms [21]. On the other hand, RT-qPCR is considered a more targeted and quantitative method, and it might show smaller fold changes because it focuses on specific regions of the transcript and might not capture all isoforms or post-transcriptional variations [22]. Nonetheless, both methods confirmed upregulation of the DEGs **(Table 1)**. HUVECs are widely utilized and well-established in vitro model to study endothelial function in cardiovascular research. They are considered a gold standard for investigating endothelial cell function and signaling under both physiological and pathological conditions. However, given the heterogeneity in endothelial cells depending on venous or arterial origin, we also validated upregulated DEGs in BRCA2-silenced and control HCAECs, which shows the similar significant up-regulation for top 10 upregulated DEGs in BRCA2-silenced HCAECs (**Table 1**). When the expression level of top downregulated DEGs were analyzed, our validation qPCR data showed similar trend for *MRC1, RAET1E, CMPK1, LRRTM2*, and *NAT8B* as all these genes were significantly downregulated in BRCA2-deficient endothelial cells **(Table 2)**. The discrepancies observed in gene expression changes between RNA sequencing and RT-qPCR could be attributed to factors such as differences in normalization strategies between the two methods, post-transcriptional regulation, or sample variability. In RNA sequencing, normalization typically adjusts for sequencing depth and gene length, while RT-qPCR normalizes against reference genes.

## Discussion

Previous studies have shown that BRCA1/2-deficient endothelial cells exhibit increased inflammation, oxidative stress, and DNA damage, ultimately leading to apoptosis and endothelial dysfunction [10, 19], all of which contribute to the development of CVDs. These observations underscore the need to further investigate the molecular pathways and targets linked to BRCA2 in endothelial biology. In this study, we performed a transcriptomic analysis of protein-coding genes to characterize gene expression changes in *BRCA2*-silenced HUVECs.

Among the top DEGs, *CEND1* (*cell cycle exit and neuronal differentiation 1*) was the most significantly upregulated gene in BRCA2-deficient endothelial cell. *CEND1* is a neuronal lineage-specific membrane protein known to coordinate cell cycle exit and differentiation [23-26]. It reduces cell proliferation through the p53/Cyclin D1/pRb signaling axis [27], and *BRCA2* silencing has previously been shown to activate p53 and p21 [19]. Thus, *CEND1* upregulation may represent a compensatory mechanism that limits proliferation and promotes cell cycle arrest in response to BRCA2 deficiency. However, because HUVECs lack neuronal differentiation machinery, this may instead predispose cells to apoptosis, consistent with earlier reports that *CEND1* overexpression induces apoptosis in non-neuronal cells such as fibroblasts [24]. *KCNAB1 (potassium voltage-gated channel subfamily A regulatory beta subunit 1)*, a gene encoding a subunit of voltage-gated potassium channels [28], is broadly expressed in both healthy and diseased human cardiac tissue [29]. Its upregulation in HUVECs exposed to postprandial hyperlipidemic serum suggests a role in the endothelial stress response [30]. Given that BRCA2 deficiency induces cellular stress, *KCNAB1* upregulation may reflect a compensatory mechanism. Notably, *KCNAB1* also promotes endothelial cell migration [31], suggesting a potential role in vascular repair. *JAKMIP2* (*Janus kinase and microtubule-interacting protein 2*), a Golgi-associated protein [32], is upregulated in contexts of DNA damage, such as after treatment with demethoxycurcumin in non-small cell lung cancer [33]. Given the DNA damage and apoptosis observed in BRCA2-deficient HUVECs exposed to DNA stressors [19], *JAKMIP2* upregulation may help maintain cell integrity under stress. *CCDC136* (*coiled-coil domain-containing protein 136*) functions as a single-pass membrane protein and is suggested to act as a tumor suppressor [34, 35]. Its upregulation has been associated with reduced angiogenesis and downregulation of Wnt/β-catenin signaling in cervical cancer [36]. Because this pathway is also critical for endothelial angiogenesis [37], *CCDC136* upregulation in *BRCA2*-silenced HUVECs may inhibit Wnt/β-catenin signaling, reducing angiogenesis and proliferation in a protective response to genomic stress. This is consistent with previous findings showing *BRCA2* knockdown suppresses angiogenesis in HUVECs [19]. *FAM83B* is an oncogene that promotes cell growth via PI3K/Akt/mTOR signaling [38]. BRCA2 silencing appears to reduce Akt activation [19], potentially triggering a feedback loop in which *FAM83B* expression increases to restore survival signaling and counteract impaired cell proliferation. *STEAP3* (*six-transmembrane epithelial antigen of prostate 3*) is a p53 transcriptional target involved in apoptosis and cell cycle regulation [39]. Its upregulation in BRCA2-deficient HUVECs likely reflects increased p53 activity [19], indicating a cellular shift toward apoptosis or altered cell cycle dynamics. As *STEAP3* is also a marker of endothelial identity [40], its elevated expression may suggest broader changes in endothelial function under stress. *IL-10RB* (*interleukin-10 receptor subunit beta*) is a cytokine receptor that suppresses immune responses and is expressed in human endothelial cells [41] [42]. BRCA2 deficiency activates the STING pathway, promoting proinflammatory cytokines such as TNF-α [15]. Upregulation of *IL-10RB* might therefore serve as a counter-regulatory mechanism to reduce inflammation and limit TNF-α-induced apoptosis. *CLMP* (*coxsackie- and adenovirus receptor-like membrane protein*) is part of the CTX family, contributing to tight junctions and leukocyte transmigration [43]. Its increased expression may reinforce endothelial barriers and limit immune cell infiltration in response to the proinflammatory state induced by BRCA2 inactivation [15, 44]. *EDIL3* (*EGF-like repeat and discoidin I-like domain-containing protein 3*) is a secreted ECM protein with angiogenic properties [45] [46, 47] and known to reduce immune cell adhesion [48, 49]. It promotes endothelial adhesion and migration [50], and its knockdown impairs retinal angiogenesis [51]. In BRCA2-deficient HUVECs, *EDIL3* upregulation may compensate for impaired angiogenesis [19] and serve as a protective mechanism against immune activation. Together, the upregulation of these genes highlights a complex cellular response to BRCA2 deficiency in HUVECs, characterized by the activation of stress adaptation pathways, modulation of proliferation, and immune regulation. These compensatory mechanisms may serve to preserve endothelial function under genomic stress but may also contribute to impaired angiogenesis and increased susceptibility to apoptosis.

The top downregulated genes; *MRC1, RAET1E, DNASE1L3, CMPK1, LRRTM2*, and *NAT8B* in BRCA2-deficient endothelial cells is likely to compromise both vascular function and cell survival. MRC1, a scavenger receptor involved in endocytosis and immune regulation, helps maintain endothelial homeostasis; its loss can impair clearance of glycoproteins and promote a pro-inflammatory environment [52]. Reduced RAET1E, a stress-induced ligand for immune recognition, may blunt endothelial stress signaling and alter interactions with natural killer (NK) cells, weakening immune surveillance [53]. Suppression of DNASE1L3, a nuclease critical for degrading extracellular DNA, may lead to accumulation of DNA debris and heightened inflammatory signaling [54]. Downregulation of CMPK1, a nucleotide kinase essential for pyrimidine metabolism, is expected to limit nucleotide pools, impairing DNA/RNA synthesis and reducing proliferative and repair capacity in already DNA-repair–deficient BRCA2-null cells [55]. LRRTM2 is best characterized as a synaptic adhesion molecule; while direct endothelial LRRTM2 studies are limited, synaptic adhesion proteins (neurexins/neuroligins/LRRTM family members) have been reported to be expressed in the vascular wall and to influence cell–cell contacts, so LRRTM2 downregulation could plausibly disrupt endothelial junction integrity and barrier stability [56]. Finally, decreased NAT8B, an ER-resident acetyltransferase (ATase1/NAT8B), could weaken endothelial stress resilience by impairing ER proteostasis and metabolic adaptation [57, 58]. Collectively, suppression of these genes amplifies the vulnerability of BRCA2-deficient endothelial cells, reducing their ability to repair, communicate, and survive under stress, ultimately predisposing the vasculature to dysfunction and injury.

Our GO analyses revealed enrichment in biological processes such as axon guidance, cholesterol biosynthesis, and regulation of cell migration and angiogenesis [59, 60]. These processes are essential for maintaining vascular integrity and function. The most enriched cellular components included plasma membrane rafts and caveolae, which are critical for endothelial cell signaling and mechanotransduction [61]. Molecular function analyses highlighted alterations in activities related to semaphorin receptor binding and phosphatase activity, indicating potential disruptions in cellular communication and signaling pathways. Moreover, KEGG pathway enrichment analyses identified significant alterations in pathways such as AGE-RAGE signaling, focal adhesion, and cGMP-PKG signaling, all of which are implicated in endothelial cell function and vascular health [62]. These molecular alterations suggest that BRCA2 deficiency impairs endothelial cell function through multiple mechanisms. Disruptions in cholesterol biosynthesis and plasma membrane organization can affect membrane fluidity and receptor function, leading to impaired signal transduction and endothelial cell responsiveness [60, 61]. Alterations in cell migration and angiogenesis pathways may hinder endothelial repair and neovascularization, compromising vascular integrity. Dysregulation of AGE-RAGE signaling and cGMP-PKG pathways can lead to increased oxidative stress and inflammation, further contributing to endothelial dysfunction [62]. Collectively, these findings underscore the critical role of BRCA2 in maintaining endothelial cell function and highlight the potential vascular implications of BRCA2 deficiency.

While our current study highlights the transcriptional impact of BRCA2 deficiency in endothelial cells, it is important to note that the associated molecular pathways may differ depending on the presence of external stressors such as hyperglycemia, hyperlipidemia, or genotoxic agents. Overall, BRCA2 deficiency induces substantial transcriptional changes that may trigger endothelial dysfunction, providing a valuable gene list and mechanistic hypotheses for future vascular biology research. Although HUVECs offer a well-established in vitro model, endothelial heterogeneity across vascular beds necessitates validation in aortic, microvascular, and macrovascular endothelial cells [63]. Additionally, the small RNA-seq sample size (n = 4 per group) limits statistical power, highlighting the need for larger studies to improve robustness, reproducibility, and generalizability of these findings.

